# HiFive: a tool suite for easy and efficient HiC and 5C data analysis

**DOI:** 10.1101/009951

**Authors:** Michael EG Sauria, Jennifer E Phillips-Cremins, Victor G Corces, James Taylor

## Abstract

The chromatin interaction assays 5C and HiC have advanced our understanding of genomic spatial organization but analysis approaches for these data are limited by usability and flexibility. The HiFive tool suite provides efficient data handling and a variety of normalization approaches for easy, fast analysis and method comparison. Integration of MPI-based parallelization allows scalability and rapid processing time. In addition to single-command analysis of an entire experiment from mapped reads to interaction values, HiFive has been integrated into the open-source, web-based platform Galaxy to connect users with computational resources and a graphical interface. HiFive is open-source software available from http://taylorlab.org/software/hifive/.

## Background

In the more than a decade since the vast majority of the human genome was first sequenced, it has become clear that sequence alone is insufficient to explain the complex gene and RNA regulatory patterns seen over time and across cell type in eukaryotes. The context of specific sequences—whether from combinations of DNA binding transcription factors (TFs) [1–3], methylation of the DNA itself [4, 5], or local histone modifications [4, 6]—is integral to how the cell utilizes each sequence element. Although we have known about the potential roles that sequentially distant but spatially proximal sequences and their binding and epigenetic contexts play in regulating expression and function, it has only been over the past decade that new sequencing-based techniques have enabled high-throughput analysis of higher-order structures of chromatin and investigation into how these structures interact amongst themselves and with other genomic elements to influence cellular function.

Several different sequencing methods for assessing chromatin interactions have been devised, all based on preferentially ligating spatially close DNA sequence fragments. These approaches include ChIA-Pet [7], tethered chromosome capture [8], and the chromatin conformation capture technologies of 3C, 4C, 5C, and HiC [9–12] (Supplemental Fig. 1). While these assays have allowed a rapid expansion of our understanding of the nature of genome structure, they also have presented some formidable challenges.

In both HiC and 5C, systematic biases resulting from the nature of the assays have been observed [13, 14], resulting in differential representation of sequences in the resulting datasets. While analyses at a larger scale are not dramatically affected by these biases due to the large number of data points being averaged over, higher-resolution approaches must first address these challenges. This is becoming more important as the resolution of experiments is increasing [15]. Several analysis methods have been described in the literature and applied to correcting biases in HiC [14–21] and 5C data [22–24]. There is still room for improving our ability to remove this systematic noise from the data and resolve finer-scale features and, perhaps more importantly, for improving the usability and reproducibility of normalization methodologies.

A second challenge posed by data from these types of assays is one of resources. Unlike other next-generation sequencing assays where even single-base resolution is limited to a few billion data points, these assays assess pairwise combinations, potentially increasing the size of the dataset by several orders of magnitude. For a three billion base-pair genome cut with a six-base restriction enzyme (RE), the number of potential interaction pairs is more than half a trillion (if considering both fragment ends) while a four-base RE can yield more than two and a half quadrillion interaction pairs. Even allowing that the vast majority of those interactions will be absent from the sequencing data, the amount of information that needs to be handled and the complexity of normalizing these data still pose a major computational hurdle, especially for investigators without access to substantial computational resources.

Here we describe HiFive, a suite of tools developed for handling both HiC and 5C data using a combination of empirically determined and probabilistic signal modeling. HiFive has performance on par or better than other available methodologies while showing superior speed and efficient memory usage through parallelization and data management strategies. In addition to providing a simple interface with no preprocessing or reformatting requirements, HiFive offers a variety of normalization approaches including versions of all commonly used algorithmic approaches allowing for straightforward optimization and method comparison within a single application. In addition to its command line interface, HiFive is also available through Galaxy, an open-source web-based platform, connecting users with computational resources and the ability to store and share their 5C and HiC analyses. All of these aspects of HiFive make it simple to use and fast, and make its analyses easily reproducible.

## The HiFive analysis suite

HiFive was designed with three goals: First, to provide a simple to use interface for flexible chromatin interaction data analysis; second, to provide well-documented support for 5C analysis; third, to improve performance over existing methodologies while reducing analysis runtimes. These are accomplished through a stand-alone program built on a Python library designed for customizable analysis and supported under the web-based platform Galaxy.

### User interface

HiFive provides three methods of use: the command line; the Internet; or as a development library. The command line interface provides users with the ability to perform analyses as a series of steps or as a single unified analysis. The only inputs that HiFive requires are a description of the genomic partitioning and interaction data, either directly as mapped reads or counts of reads associated with the partitioned genome (e.g. fragment pairs and their observed reads). HiFive handles all other formatting and data processing. In addition, HiFive has been bundled as a set of tools available through Galaxy (Fig. 1). This not only provides support with computational resources but also ensures simple installation of all prerequisite libraries and packages. HiFive was also created to allow custom creation of analysis methods as a development library for chromatin interaction analysis through extensive documentation and an efficient data-handling framework.

**Figure 1.**
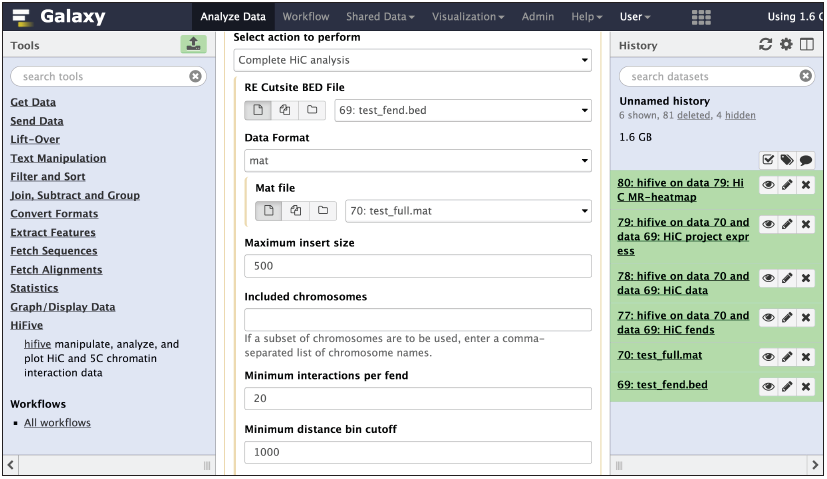
HiFive’s tool interface through Galaxy. HiFive tools are available through the Galaxy toolshed, providing a graphical interface and showing tool option inter-dependencies.

### Organization of HiFive

At its core, HiFive is a series of hierarchical data structures building from general to specific information. There are four primary file types that HiFive creates, all relying on the Numpy scientific computing Python package for efficient data arrays and fast data access. These structures include genomic partitioning, observed interactions, distance-dependence relationship and normalization parameters, and heatmaps of observed and expected interactions. By separating these attributes, many datasets can utilize the same genomic partitioning and multiple analyses can be run using the same interaction data without the need to reload or process information.

### Data processing and filtering

In order to process 5C or HiC data, the first step after mapping is converting the data into a form compatible with the desired workflow. HiFive appears to be nearly alone in its ability to handle mapped data without additional processing (the only exception is HiCLib [17]). Reads can be read directly from numerous BAM-formatted files and this may be done as an independent step or within the integrated one-step analysis. HiCLib also possesses the ability to input data directly from mapped read files. In all other cases, reads need to be converted to other formats. HiCPipe [14] provides scripts for some but not all of these processes, while HiCNorm [16] relies on pre-binned reads. In all cases aside from HiFive, a workflow is required to move from mapped reads to normalization.

Filtering is accomplished in two phases, during the initial processing of reads and during project creation (Supplemental Figs. 2 & 3). The first phase limits data to acceptable paired-end combinations. For 5C data, this means reads mapping to fragments probed with opposite-orientation primers. HiC data uses two criteria, total insert size (a user-specified parameter) and orientation/fragment relationship filtering. In the latter case, reads originating from non-adjacent fragments or from adjacent fragments and in the same orientation are allowed, similar to Jin F, *et al.* [19] (Supplemental Fig. 4). The second phase, common to both 5C and HiC data, is an iterative filtering based on numbers of interactions per fragment or fragment end (fend). Briefly, total numbers of interactions for each fragment are calculated and fragments with insufficient numbers of interaction partners are removed along with all of their interactions. This is repeated until all fragments interact with a sufficient number of other non-filtered fragments. This filtering is crucial for any fragment or fend-specific normalization scheme to ensure sufficient interdependency between interaction subsets to avoid convergence issues.

### Distance-dependence signal estimation

One feature of HiFive that is notably absent from nearly all other available analysis software is the ability to incorporate the effects of sequential distance into the normalization. One exception to this is HiTC [21], which uses a loess regression to approximate the distance-dependence relationship of 5C data to genomic distance. This method does not, however, allow for any other normalization of 5C data. Another is Fit-Hi-C [25], although this software assigns confidence estimates to mid-range contact bins rather than normalizing entire datasets. This feature is of particular importance for analysis of short-range interactions such as this in 5C data, or for making use of counts data rather than a binary observed/unobserved indicator. For 5C data, HiFive uses a linear regression to estimate parameters for the relationship between the log-distance and log-counts (Supplemental Fig. 5). HiC data requires a more nuanced approximation because of the amount of data involved and the non-linear relationship over the range of distances queried. To achieve this, HiFive uses a linear piecewise function to approximate the distance-dependent portion of the HiC signal, similar but distinct from that used by Fit-Hi-C. HiFive partitions the total range of interactions into equally sized log-transformed distance bins with the exception of the smallest bin, whose upper bound is specified by the user. Mean counts and log-transformed distances are calculated for each bin and a line is used to connect each set of adjacent bin points (Supplemental Fig. 6). For distances extending past the first and last bins, the line segment is simply extended from the last pair of bins on either end. Simultaneously, a similar distance-dependence function is constructed using a binary indicator of observed/unobserved instead of read counts for each fend pair. All distances are measured between fragment or fend midpoints.

### HiFive normalization algorithms

HiFive offers three different normalization approaches. These include a combinatorial probability model based on HiCPipe’s algorithm called ‘Binning’, a modified matrix-balancing approach called ‘Express’, and a multiplicative probability model called ‘Probability’. In the Binning algorithm, learning is accomplished in an iterative fashion by maximizing each set of characteristic bin combinations independently each round using the Broyden–Fletcher–Goldfarb–Shanno algorithm for maximum likelihood estimation.

The Express algorithm is a generalized version of matrix balancing. While it can use the Knight-Ruiz algorithm [26] for extremely fast standard matrix balancing (ExpressKR), the Express algorithm also has the ability to take into account differing numbers of possible interactions and find corrections weighted by these numbers of interactions. The set of valid interactions is defined as set *A,* interactions whose fends have both passed the above-described filtering process and whose inter-fend distance falls within user-specified limits. In addition, because counts are log-transformed for 5C normalization, only non-zero interactions are included in set *A*. For each interaction *c* between fends or fragments *i* and *j* for HiC and 5C, respectively in the set of valid interactions *A*, correction parameter *f_i_* is updated as in (1) for HiC and (2) for 5C.

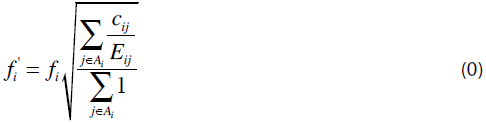

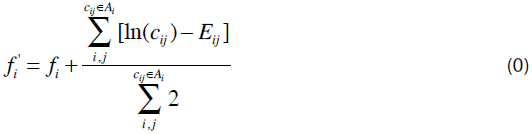

The expected value of each HiC interaction is simply product of the exponent of the expected distance-dependent signal *D(i,j)* and the fend corrections (3).

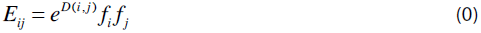

5C interactions have expected values that correspond to the log-transformed count and are the sum of each signal component (4).

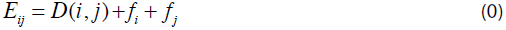

By scaling the row sums based on number of interactions, the weighted matrix balancing allows exclusion of interactions based on interaction size criteria without skewing correction factors due to non-uniform restriction site distribution, position along the chromosome, or filtered fragments or fends due to read coverage. Because it can incorporate the distance-dependent signal, the Express algorithm can operate on counts data unlike most other matrix balancing approaches, although it also can be performed on binary data (observed vs. unobserved) or log-transformed counts for HiC and 5C, respectively. This algorithm allows for adjustment of counts based on the estimated distance-dependence signal prior to normalization in both weighted (1 & 2) and unweighted (Knight-Ruiz) versions.

The multiplicative Probability algorithm models the data assuming some probability distribution with a prior equal to the estimated distance-dependent signal. HiC data can be modeled either with a Poisson or binomial distribution. In the case of the binomial distribution, counts are transformed into a binary indicator of observed/unobserved and the distance-dependence approximation function is based on this same binary data. 5C data are modeled using a lognormal distribution. In both cases only counts in the set of reads *A* (described above) are modeled.

For both the Express and Probability algorithms, a backtracking-line gradient descent approach is used for learning correction parameters. This allows the learning rate *r* to be updated each iteration *t* to satisfy the Armijo criteria (5) based on the cost C, ensuring that parameter updates are appropriate.

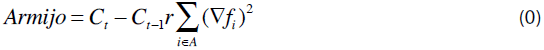

### Filtering interactions by interaction size

Chromatin topology is organized around highly reproducible regions of frequent local interactions termed ‘topological domains’ [27]. Within these structures it has been observed that specific features can influence the frequency of interactions in a biased and differential way up- and downstream of them, such as transcript start sites and CTCF bound sites[14]. In order to account for systematic noise and bias without confounding normalization efforts with meaningful biological-relevant structures, HiFive allows filtering out of interactions using interaction size cutoffs. In order to assess the effects of filtering out shorter-sized interactions, we analyzed data both with and without a lower interaction distance cutoff. For HiC data we analyzed two mouse ESC datasets with no lower limit and with a lower distance limit of 500 Kb using each of the described normalization algorithms. This size was chosen to eliminate all but the weakest interaction effects observed for TSSs and CTCF-bound sites [28]. HiC normalization performance was assessed using the inter-dataset correlations. For 5C data, there is a much smaller range of interactions. In order to handle this, we set a lower interaction size cutoff of 50 Kb. 5C normalization performance was assessed as the correlation between 5C data and HiC data of the same cell type [27] and binned based on probed 5C fragments to create identically partitioned sets of interactions and normalized using HiFive’s Probability algorithm.

The differences in HiC algorithm performances with and without the lower interaction size cutoff were varied, although the largest effects were seen when data were binned in 10 and 50 Kb bins for intra-chromosomal interactions and for overall inter-chromosomal interactions (Supplemental Fig. 7). Overall, excluding short-range interactions made little difference for the Express algorithm but did improve the performance of the Probability and Binning algorithms. The 5C algorithms showed an opposite result, with almost universal decrease in performance when short-range interactions are excluded (Supplemental Fig. 8). As a result, learning HiC normalization parameters using HiFive algorithms was performed excluding interactions shorter than 500 Kb and 5C analyses were performed using all interaction sizes. All analyses subsequent to normalization (e.g. dataset correlations) were performed across all interactions.

### Analyzing 5C data

To date, limited work has focused on processing of 5C data to remove technical biases [22–24, 29]. Of that, none has been formalized in published analysis software. In order to assess HiFive’s performance in normalizing 5C data, we used two different 5C mouse embryonic stem cell datasets [23, 24] and found correlations to HiC data of the same cell type [27] and binned based on probed 5C fragments to create identically partitioned sets of interactions (Fig. 2, Supplemental Figs. 8 & 10). HiC interactions were normalized using either HiFive’s probability algorithm (Fig. 2) or HiCPipe (Supplemental Fig. 10) and heatmaps were dynamically binned to account for sparse coverage (see Supplemental Methods: 5C-HiC data correlations).

Correlations were found between all non-zero pairs of bins (fragment level resolution) following log-transformation. All of HiFive’s 5C algorithms showed an improved correlation with HiC data compared to raw data, regardless of HiC normalization approach. The Binning algorithm showed the least improvement, likely due to the limits on the number of bins into which features could be partitioned and characteristics missing from the model, such as primer melting temperature. The standard matrix-balancing approach (ExpressKR) showed good improvement, although not quite as good as the Express and Probability algorithms. All of these normalizations were accomplished in less than a minute proceeding from a bed file and a counts file to heatmaps.

**Figure 2.**
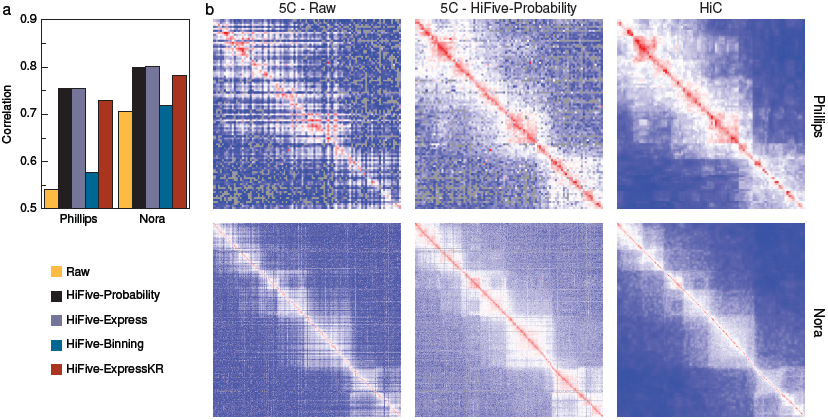
5C analysis performance. HiFive normalization of 5C data and their correlation to corresponding HiC data. a) Correlation of 5C data (intra-regional only) with the same cell type and bin-coordinates in HiC data, normalized using HiFive’s probability algorithm for two different datasets and using each of HiFive’s algorithms. b) Heatmaps for a select region from each dataset, un-normalized, normalized using HiFive’s probability algorithm, and the corresponding HiC data, normalized and dynamically binned.

## HiC analysis software comparison

Several algorithms have been proposed to handle interaction data normalization (Table 1). These analysis approaches can be divided into two varieties, probabilistic and matrix balancing. The probabilistic approach is further divided into combinatorial and multiplicative corrections. The combinatorial probability model is implemented in HiCPipe [14] and remains one of the most popular approaches. This approach uses one or more restriction fend characteristics partitioned into ranges of values and iteratively learns correction values for each combination of ranges based on a binomial distribution of observed versus unobserved fend interactions. A multiplicative modeling approach is used in the analysis software HiCNorm [16]. HiCNorm uses a Poisson regression model using binned counts instead of binary output and assuming that biases from fend characteristics are multiplicative between bin combinations. A different multiplicative approach is matrix balancing, which finds a value for each row/column of a symmetric matrix (or in this case, heatmap) such that after multiplication of each matrix value by its associated row and column values, the sum of each row and column is one. This has been described with at least four different implementations in the literature [15, 17, 20, 30] although only two software packages making use of it has been published (HiCLib [17], now included in the R package HiTC [21] and Hi-Corrector [20]). For this paper, we chose to use our own implementation of the algorithm described by Knight and Ruiz [26] for comparison due to speed and ease of use considerations.

**Table 1.** A comparison of HiC Analysis software algorithms and features.

| Method | Normalization algorithm | Can account for distance | Fend-level resolution | Parallelizable |
| --- | --- | --- | --- | --- |
| HiFive | Matrix Balancing | X | X | X |
|  | Binning | X | X | X |
|  | Probability | X | X | X |
| HiCPipe | Binning |  | X | X |
| HiCNorm | Poisson Regression |  |  |  |
| HiCLib | Matrix Balancing |  | X |  |
| HiTC* | Poisson Regression |  |  |  |
|  | Matrix Balancing |  | X |  |
| HOMER | Read Coverage | X |  | X |
| Hi-Corrector | Matrix Balancing |  | X | X |
| *This method is an R-based implementation of HiCLib's and HiCNorm's normalization approaches. |  |  |  |  |

### Method performances

To assess HiC analysis method performances we used two different pairs of HiC datasets [15, 27, 31], finding interaction correlations across different restriction digests of the same cell type genomes. The Dixon et al. data [27] was produced using mouse ES cells digested with either HindIII or NcoI, yielding approximately 4 kilobase (Kb) fragments. The Selvaraj et al. data [31] was produced from human GM12878 cells using HindIII, while the Rao et al. data [15] was produced from human GM12878 cells using the 4 base pair (bp) restriction enzyme MboI, producing approximately 250 bp fragments. This allowed assessment of method performance and data handling across a range of experimental resolutions. Correlations were calculated for ten mutually exclusive intra-chromosomal (cis) interaction ranges and across all cis interactions simultaneously for four binning resolutions. Correlations were also calculated for inter-chromosomal interactions for two resolutions.

HiC analysis methods showed varied performances across interaction size ranges, resolutions, and datasets for intra-chromosomal interactions (Fig. 3a & b). For small interaction sizes, HiFive’s Probability and Express algorithms performed consistently well regardless of resolution. At longer interaction distances the Express algorithm typically outperformed the Probability algorithm. HiCNorm showed a nearly opposite performance with poorer interdataset correlations for shorter-range interactions but higher correlations at longer ranges, relative to other methods. HiCPipe’s performance appeared to depend on binning resolution. At higher resolutions (≤ 50 Kb), HiCPipe performs worse than the majority of methods. However at lower resolutions it tends to outperform other methods, regardless of interaction size range. HiFive’s Binning algorithm had a more consistent performance around the middle of all of the methods across all binning resolutions, with the exception of the 1Mb resolution for the human data where it performed the worst. Standard matrix balancing consistently performed at or near the bottom of the group regardless of the interaction size range or resolution.

Correlations across all intra-chromosomal interactions showed much more consistency between analysis methodologies (Fig. 3c & d). This is primarily due to the fact that the main driver of inter-dataset correlation, the interaction distance-counts relationship, was present in all of the analyzed data. HiFive’s Probability and Express algorithms were again top performers across almost every intra-chromosomal comparison, although the Probability algorithm showed a decreasing advantage with decreasing binning resolution. HiCNorm, HiCPipe, matrix balancing, and HiFive’s Binning algorithm were highly consistent in terms of performance for the mouse datasets. For the human inter-dataset correlations HiCPipe and matrix balancing showed a slightly better performance than average while HiCNorm faired worse. HiFive’s Express algorithm was still the top performer.

**Figure 3.**
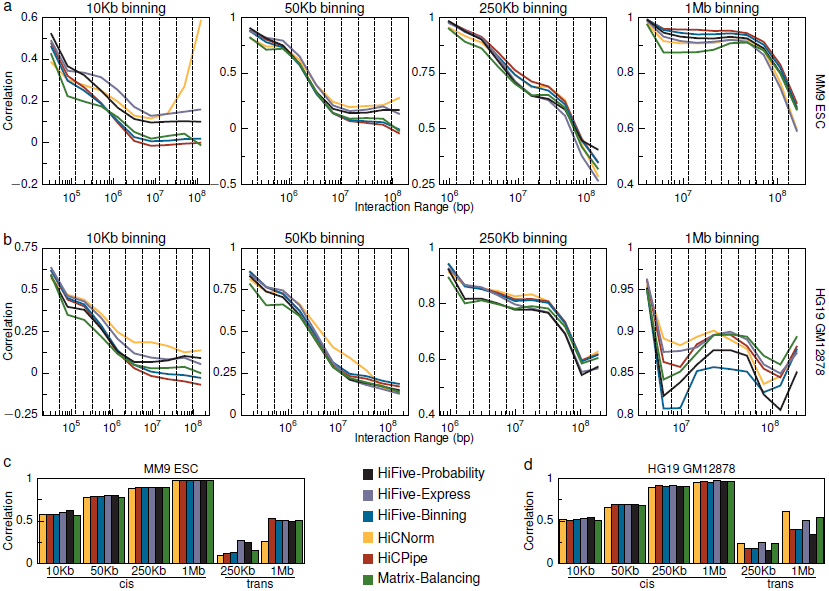
HiC method comparison. Interaction correlations between datasets created with different restriction enzymes for multiple normalization schemes across different binning resolutions. Two datasets are shown, mouse and human. Each mouse dataset was produced using a six-base restriction enzyme. The human datasets were mixed, one produced with a six-base cutter and the other with a four-base cutter. a) Data were normalized using several approaches and compared for correlation between two mouse HiC datasets. Interactions were broken down into ten groups of non-overlapping cis interaction ranges for four resolution levels. b) Correlations for ten different non-overlapping cis interaction ranges at each resolution for each analysis approach. c) Overall mouse dataset correlations for each resolution for intra-chromosomal (cis) and inter-chromosomal (trans) interactions. d) Overall human dataset correlations for each resolution for intra-chromosomal (cis) and interchromosomal (trans) interactions.

Inter-chromosomal datasets showed a wider range of performances and were strongly dependent on which datasets were being analyzed (Fig. 3c & d). For mouse interchromosomal interactions, HiFive’s Probability and Express algorithms performed much better than other methods at the 250 Kb binning resolution, but consistent with other methods at the 1 Mb resolution. HiCNorm showed worse performance at both bin sizes for the mouse datasets. HiCPipe showed the best performance at the 1 Mb resolution, slightly above other methods, but the second worst performance at the 250 Kb resolution. Results for the human datasets were more consistent across resolutions. HiCNorm, HiFive’s Express algorithm, and matrix balancing performed best in both cases with Express doing slightly better at the 250 Kb resolution and HiCNorm at the 1 Mb resolution. The remaining methods showed similar performance to each other, although HiFive’s Probability algorithm performed slightly worse than HiFive’s Binning algorithm and HiCPipe.

The inconsistency between results for cis and trans interactions suggests that no approach is ideal for both types of interactions. To further explore this we looked at the effects of pseudocounts in the Binning/HiCPipe normalization scheme and the effects of distance-dependence on normalization. Pseudo-counts are values added to both expected and observed reads to mitigate the impact of stochastic effects. HiCPipe showed a stronger performance compared to HiFive’s Binning algorithm at longer ranges and at larger bin sizes. We determined that the primary difference was the inclusion of pseudo-counts in all feature bins prior to normalization. By progressively adding counts, we found that cis interaction correlations decreased at shorter interaction ranges and overall although the correlations increased at longer ranges and for trans interactions (Supplemental Fig. 11).

We also performed parallel analyses using our weighted matrix balancing algorithm, Express, with and without the estimated distance-dependence signal removed prior to normalization (Supplemental Fig. 12). This showed a similar effect to the addition of pseudo-counts, such that leaving the distance-signal relationship intact resulted in stronger long-range interaction correlations in larger bin sizes, stronger 1 Mb binned trans correlations, and poorer overall cis interaction correlations across all bin sizes.

### Computational requirements

In order to determine the computational requirements of each analysis method, we ran each analysis on an abbreviated dataset consisting of a single chromosome of cis interactions from the mouse NcoI dataset starting from loading data through producing a 10 Kb heatmap. All normalizations were run using a single processor and publicly available scripts/programs. The exception to this was binning the counts and fragment feature data for HiCNorm. No script was provided for this step so one was written in R to supplement HiCNorm’s functionality.

Runtimes varied greatly between normalization methods, ranging from less than 7 minutes to approximately 12.5 hours (Fig. 4). With the exception of HiFive’s Probability algorithm, HiFive performed better in terms of runtime than all other algorithms. HiCPipe and HiCNorm both showed long runtimes at least an order of magnitude above other methods. The slowest approach, though, was HiFive’s Probability algorithm. This was due to its modeling of every interaction combination across the chromosome. HiFive’s implementation of the Knight-Ruiz matrix balancing algorithm, ExpressKR, showed a dramatically faster runtime than any other approach. This was the result of HiFive’s fast data loading and efficient heatmapping without the need for distance-dependence parameter calculations.

**Figure 4.**
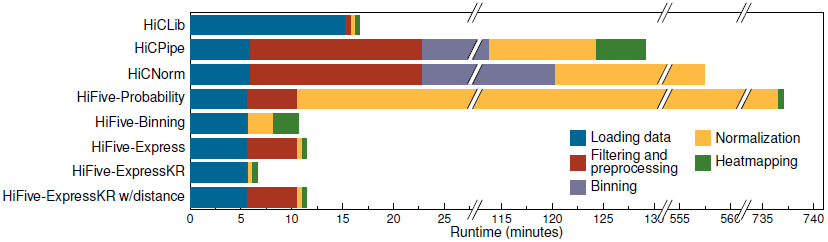
Running time for HiC analysis methods. For each method, the runtime in minutes is partitioned into the time required for each stage of the processes. All times were determined running methods on an abbreviated dataset of chromosome 1 for the mouse HindIII dataset using a single processor. Note that because of several extremely long runtimes, the graph includes multiple splits.

### Scalability

Because of the ever-increasing resolution of experiments and the corresponding size of interaction datasets, scalability is a primary concern for HiC data analysis. Although we compared methods on an even playing field, this does not reflect the complete performance picture when considering finer-scale resolution, processing a complete genome interaction dataset, and more available computational resources.

There are two approaches to determining analysis resolution, prior to or after normalization. Of the methods presented, only HiCNorm determines the resolution of analysis prior to normalization. While it performs well, this means that the processing time and memory requirements scale exponentially with resolution. We were unable to perform any analyses at resolutions for bin sizes smaller than 10 Kb using this approach. The remaining methods all find correction values for individual fends, meaning that corrections are performed prior to binning interactions.

The increase in dataset size, either due to genome size itself or a finer-scale partitioning of the genome, can be offset by employing more processing power by means of parallelization. HiCLib and HiCNorm do not appear to have any such capability. HiCPipe does have the ability to parallelize calculation of model bin sizes prior to normalization and calculations for heatmap production, although a single processor performs all of the actual normalization calculations. HiFive, on the other hand, has the ability to run in parallel for nearly all phases of an analysis. The two exceptions are loading the initial data and filtering reads, although the latter is very fast already. All normalization algorithms, including the Knight-Ruiz algorithm implemented in HiFive, have been parallelized for HiC analysis using MPI. The parallelization is implemented in such a way that the additional memory overhead for each new process is minimal.

## Conclusions

HiC analysis remains a challenging subject, as demonstrated by the varied performances across all methodologies discussed here. No single approach appears to be ideally suited for all cases, suggesting that the experimental goal should drive the choice of analysis software. It is unclear how best to assess HiC normalization performance as there is no ‘gold standard’ for determining the quality of a HiC dataset or how well systematic noise has been accounted for during an analysis. As seen in the differences in correlation between mouse and human datasets (Fig. 3), factors such as restriction fragment size distributions, cut site density, sequencing depth, and HiC protocol can dramatically impact the similarity of resulting datasets. Further, in order to detect biologically relevant features against the background of the distance-signal relationship, the data needs to be transformed, typically using a log-transformation. This skews the resulting comparison by ignoring interactions for which no reads have been observed, an increasing problem as binning size decreases or interaction size increases. At longer ranges, non-zero bins are sparse and dominated by macro features (such as A-B compartments), a situation that can result in increasing correlations (Fig. 3a & b). Two observations suggest this is not an artifact. First, the long-range interaction correlation increase is seen in the human but not mice data, reflecting differences in genome organization. Second, the correlation increases are seen across all methodologies and algorithms.

Normalization software attempts to account for many of these confounding factors and allow direct comparison between datasets produced by different labs, protocols, and even across species although what can reasonably be expected in terms of this normalization process is unclear. This question depends on many factors and we may not have sufficient understanding of chromatin architecture variability across a cell population to answer it accurately. The resolution (bin size), similarity of datasets in terms of sequencing depth, restriction fragment size distributions, and protocol, as well as cell population size and population similarity from which the HiC libraries were made will all influence the correlation. At a low resolution, say 1Mb, we should expect nearly a perfect correlation. However, at much higher resolution differences in mappability and RE cut-site frequency will strongly influence the correlation. Further, we need to consider the distance dependence of the signal as this is the strongest driver of the correlation and can give a false impression of comparability between datasets.

To address these normalization challenges, we have create HiFive, an easy-to-use, fast, and efficient framework for working with a variety of chromatin conformation data types. Because of the modular storage scheme, re-analysis and downstream analysis is made easier without additional storage or processing requirements. We have included several different normalization approaches and made nearly all aspects of each algorithm adjustable, allowing users to tune parameters for a wide range of analysis goals. HiFive is parallelized via MPI, making it highly scalable for nearly every step of HiC data processing.

For 5C data, HiFive is the only analysis program available for normalization and allows easy management of 5C data. We have demonstrated that 5C data normalizations performed by HiFive greatly improve consistency between 5C data and corresponding HiC data across multiple datasets.

We have also shown HiFive’s performance in handling HiC data. HiFive is consistently performing at or above other available methods as measured by inter-dataset correlations for cis interactions. In addition, we have demonstrated that HiFive is tunable to achieve superior trans performance if desired, albeit at the expense of performance across cis interactions. HiFive has also proved capable of handling very high-resolution data, making it useful for the next generation of HiC experimental data.

In terms of performance considerations, our analysis suggests that, out of all of the methods considered, the balance between speed and accuracy is best achieved by HiFive-Express or HiFive-ExpressKR. This appears to be true regardless of resolution or dataset size. In order to get this performance, it is crucial to use the distance-dependence adjustment prior to normalizing, necessitating the need to pre-calculate the distance-dependence function. Because this requires iterating over every possible interaction, using multiple processors is highly recommended. If not possible, HiFive-ExpressKR without distance correction is a robust fallback method. If computational resources are not a limiting factor, we recommend HiFive-Probability. With approximately 100 CPUs, the high-resolution human data was processed in about a day. At fine-scale binning, this approach yields the best results of all methods.

While HiFive allows for superior normalization of data compared to other available software under many conditions, it also provides users with alternative options for fast analysis with minimal computational requirements at only a slight accuracy cost, opening high-resolution HiC and 5C analysis to a much larger portion of the scientific community. HiFive is available at http://taylorlab.org/software/hifive/. Source code is provided under an MIT license and at https://github.com/bxlab/hifive or installed using pip from http://pypi.python.org.

## List of abbreviations used

RE: restriction enzyme; fend: fragment-end; Kb: kilobase; bp: base pair; cis: intra-chromosomal; trans: inter-chromosomal; Mb: megabase; 3C: chromosome conformation capture; 5C: 3C carbon copy; MPI: message passing interface; CPU: central processing unit.

## Authors’ contributions

MEGS, JEPC, VGC, and JT conceived the project and developed feature requirements. JEPC made significant contributions to the design of the 5C tools. MEGS developed all algorithms, designed and wrote all software, and wrote the manuscript. JT contributed to the manuscript and supported the project. All authors read and approved the final manuscript.

## Additional data files

The following additional data files are available with the online version of the paper. Additional data file 1 contains a detailed methods section, a table listing the sources of datasets used, and Supplemental figures. Additional data file 2 contains a tar archive containing the HiFive software library. Additional data file 3 contains a tar archive containing all scripts using to generate the data, analyses, and figures presented in this paper. Additional data file 4 contains the software documentation.

## Acknowledgements

Research reported in this publication was supported by the National Institutes of Health under awards R01GM035463 to VC and R01DK065806 to JT and by American Recovery and Reinvestment Act (ARRA) funds through grant number RC2HG005542 to JT. The content is solely the responsibility of the authors and does not necessarily represent the official views of the National Institutes of Health.

